# The VPS9-family GEF VINE activates Ypt10 in a late endosomal Rab cascade

**DOI:** 10.64898/2026.07.13.738292

**Authors:** Mia S. Frier, Michael Davey, Elizabeth Conibear

## Abstract

Rab GTPase cascades drive endosomal membrane maturation by sequentially activating and inactivating Rab proteins. These transitions in Rab signaling require the coordinated actions of guanine nucleotide exchange factors (GEFs) and GTPase-activating proteins (GAPs). The yeast VINE complex is an endosomal VPS9-family GEF that stimulates a GAP to inactivate the Rab5 homolog Vps21, suggesting a role for VINE in coordinating Rab transitions. Here we report that VINE acts through its catalytic GEF domain to promote signaling by the Rab5-related GTPase Ypt10 and establish a pool of Ypt10 at late endosomes. Ypt10 activation occurs downstream of Vps21 activity, placing Ypt10 within a late endosomal Rab cascade. Genome-wide protein proximity screens revealed a VINE-dependent interaction between Ypt10 and the GEF Mon1-Ccz1. Our data suggest that VINE and Ypt10 regulate late endosomal recruitment of Mon1-Ccz1 to enhance the activation of its substrate, the Rab7 homolog Ypt7. Together, these findings define a Vps21-VINE-Ypt10 regulatory module that adds a layer of control within the late endosomal Vps21-to-Ypt7 cascade and establish VINE as a dual Rab regulator. Through opposing activities on Vps21 and Ypt10, VINE may couple Rab5 inactivation to Mon1-Ccz1 recruitment to provide more precise control of degradative protein traffic to the vacuole.

**Significance statement:** Four Rab5-family GTPases direct protein sorting and membrane maturation in the yeast endolysosomal system, yet their individual functions, and the role of the little-studied Rab Ypt10, are unclear.

Using genome-wide proximity screens, we find that the GEF complex VINE establishes a pool of Ypt10 at late endosomes downstream of Vps21, where Ypt10 recruits Mon1-Ccz1, the activator of the Rab7 homolog Ypt7.

Because VINE also drives GAP-mediated suppression of Vps21, our results suggest it acts as a dual Rab regulator, coupling Vps21 inactivation to Ypt10 activation to fine-tune the endosomal Rab cascade.

## Introduction

Rab GTPases control cargo sorting, vesicle biogenesis, and membrane fusion events that together direct membrane trafficking between organelles (Pfeffer, 2017). Endosomal Rab GTPases in particular regulate the trafficking that maintains the plasma membrane and Golgi proteomes and delivers degradative cargo to the vacuole or lysosome (Langemeyer et al., 2018). To coordinate this sorting and trafficking, positive and negative Rab regulators are targeted to specific sites on endosomal membranes, where they control Rab signaling. For example, membrane coats in the endolysosomal system recruit guanine nucleotide exchange factors (GEFs), which catalyze nucleotide exchange to locally convert their target Rab GTPases into an active, GTP-bound form that promotes cargo sorting and membrane fusion (Hesketh et al., 2014; Bean et al., 2015; Antón-Plágaro et al., 2025). GTPase-activating proteins (GAPs) catalyze GTP hydrolysis to return the Rab to an inactive, GDP-bound form that cannot bind effectors, thereby dissipating the Rab signaling domain (Haas et al., 2005; Lachmann et al., 2012; Nickerson et al., 2012). The combined activity of GEFs and GAPs drives transitions in Rab signaling that are important for ongoing transport (Rivera-Molina and Novick, 2009).

Multiple GTPases in the Rab5 family recruit distinct sets of effectors and cooperate to establish endosomal membrane identity and function (Bucci et al., 1995; Chen et al., 2009; Del Olmo et al., 2019; Duarte et al., 2022). VPS9-family GEFs are similarly diverse, and each appears to play a distinct role in linking endosomal Rab activation to specific processes, such as cargo sorting into degradative or recycling pathways (Carney et al., 2006; Lee et al., 2006; Nesterova et al., 2026). Mechanisms that selectively target Rab GEFs and GAPs to endosomal membranes could regulate Rab signaling not only at stable membrane subdomains but also during dynamic processes such as membrane fusion or coat disassembly (Zhang et al., 2014; Jia et al., 2016; Law et al., 2017). Understanding the contribution of each VPS9-family GEF will therefore depend on identifying its Rab GTPase substrates and determining how Rab-GEF modules are activated in the correct spatial and temporal contexts to coordinate endosomal trafficking.

Within Rab cascades, the timed recruitment of GEFs and GAPs sequentially organizes signaling by two different Rab proteins (Novick, 2016). At the maturing endosomal membrane, for example, Rab5 recruits the GEF Mon1-Ccz1, which activates Rab7 (Rink et al., 2005; Nordmann et al., 2010). Because Rab5 inactivation is in turn promoted by various components of the Rab7 signaling network, this cascade converts a Rab5 signaling domain into a Rab7 domain prior to lysosomal delivery (Chotard et al., 2010; Poteryaev et al., 2010; Rana et al., 2015; Tu et al., 2022). Defining the mechanistic details of this reciprocal regulation offers insight into how transitions in Rab signaling are precisely coordinated with cargo trafficking and membrane maturation.

In budding yeast, endosomal membranes are populated by four Rab GTPases that carry out shared and specific functions (Singer-Krüger et al., 1994; Nesterova et al., 2026). The most abundant Rab5-like GTPase in yeast is Vps21, which has unique functions in membrane tethering and autophagy (Russell et al., 2012; Cabrera et al., 2013; Chen et al., 2014). The stress-induced Vps21 paralog Ypt53 performs similar functions (Schmidt et al., 2017). Ypt52, the second most highly expressed yeast endosomal Rab, is reported to have unique interactions with both regulators and effectors (Liu et al., 2011; Duarte et al., 2022). Finally, while effector binding by the fourth endosomal Rab, Ypt10, has been described *in vitro*, its *in vivo* functions remain largely unexplored (Louvet et al., 1999; Langemeyer et al., 2020; Nesterova et al., 2026).

Two VPS9-family GEFs, Vps9 and Muk1, play partially redundant roles in promoting signaling by these Rab GTPases (Cabrera et al., 2013; Paulsel et al., 2013; Locke and Thorner, 2018). In contrast, the regulatory roles of the related GEF Vrl1 are less well understood, because a widely used laboratory yeast lineage carries a mutation in the *VRL1* gene that prevents expression of full-length Vrl1 (Bean et al., 2015). Vrl1 is the GEF subunit of the heterodimeric VPS9 GEF-interacting sorting nexin (VINE) complex, which also contains the sorting nexin- (SNX-) Bin-amphiphysin-Rvs (BAR) adaptor Vin1 (Shortill et al., 2022). VINE localizes to endosomes where it cooperates with the phosphatase Glc7 to promote GAP-mediated inactivation of the major yeast Rab5 homolog, Vps21 (Frier et al., 2026). This pathway selectively opposes Vps21 signaling without affecting Ypt52 activity. Because it both possesses a GEF domain and promotes GAP activity, VINE has the potential to coordinate positive and negative Rab regulation during endosomal trafficking or maturation. If VINE activates a second GTPase while promoting Vps21 inactivation, it could convert a Vps21 signaling domain into one defined by a different active Rab. Further study of VINE could therefore reveal which regulatory events coincide with Vps21 inactivation and how such coordinated Rab regulation supports endosomal membrane dynamics.

Here we identify VINE as a positive regulator of the endosomal Rab5-related GTPase Ypt10. We show that downstream of Vps21 activity, VINE acts through its catalytic GEF domain to establish Ypt10 signaling at late endosomes. Our data indicate that this late endosomal pool of Ypt10 helps to recruit the Mon1-Ccz1 complex, which activates the Rab7 homolog Ypt7. Together, our findings establish VINE as a dual Rab regulator that fine-tunes the late endosomal Rab cascade through both positive and negative Rab regulation.

## Results

To understand the role of VINE in endosomal Rab regulation, we completed a genome-wide screen to describe the local molecular environment and identify close functional partners. We previously used the split-dihydrofolate reductase (DHFR) method to catalogue the proximity interactome of the VINE subunit Vrl1 and identify its binding partner, Vin1 (Tarassov et al., 2008; Shortill et al., 2022). Because the C-terminal DHFR^C^ tags used in this screen can interfere with membrane targeting and function, this initial Vrl1 interactome likely lacks important interactors such as Rab GTPases and SNAREs. To capture a more complete list of Vrl1 proximity interactors, we recently developed a Nanobody-Adapted Proximity Assay (NAPA), which is a modified split-DHFR assay in which expression of a GFP-binding nanobody (Nb) effectively converts a collection of N-terminally GFP-tagged proteins (Weill et al., 2018) into DHFR^C^ preys (Figure 1A; Frier et al., 2026).

**Figure 1.**
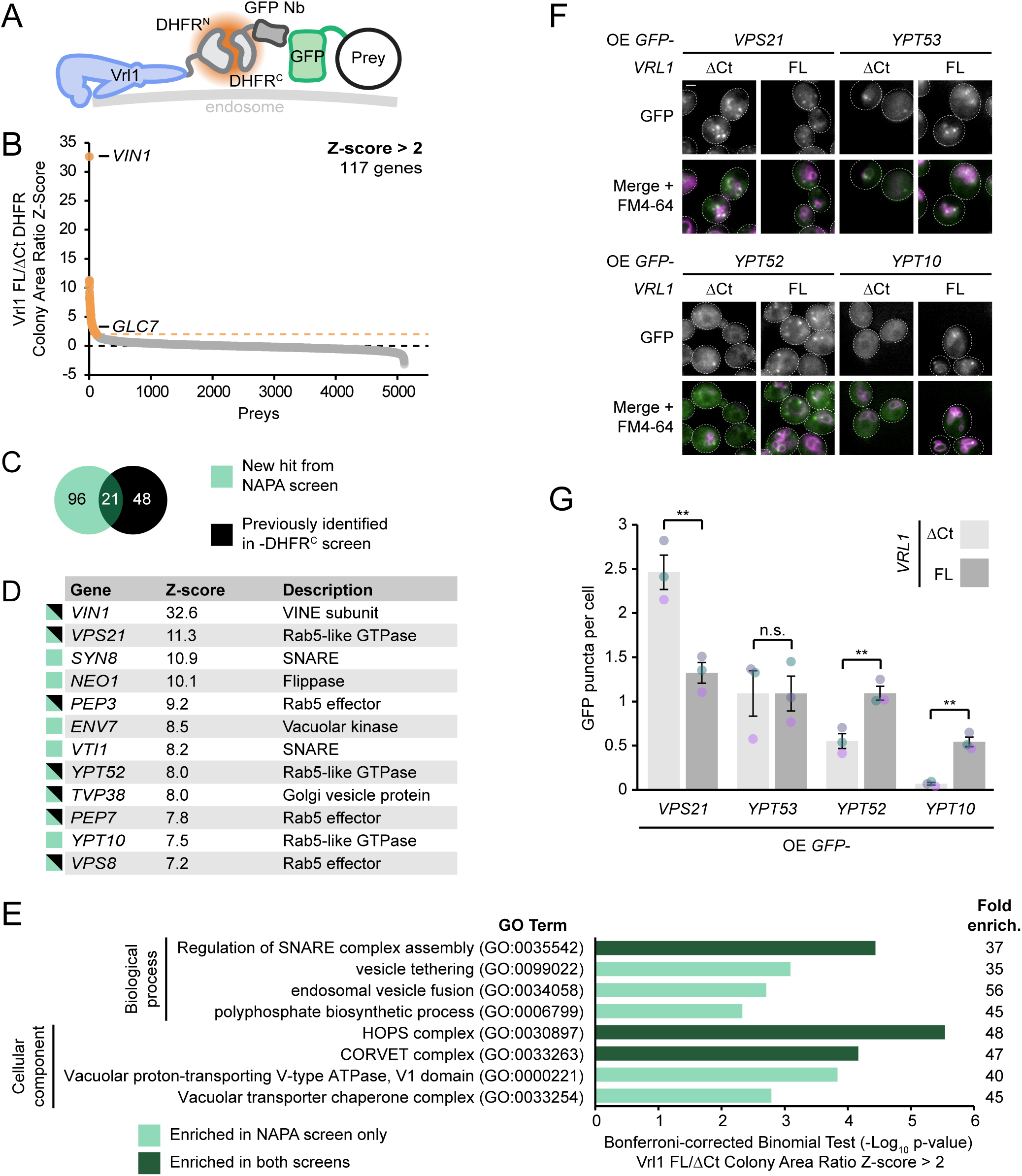
A nanobody-adapted proximity screen for VINE interactors identifies the Rab GTPase Ypt10. (A) Schematic of the nanobody-adapted proximity assay (NAPA). The Vrl1 bait is tagged at its N-terminus with the N-terminal fragment of DHFR. Prey proteins are GFP-tagged at their N-terminus, and a GFP-binding nanobody (Nb) targets the C-terminal fragment of DHFR to the prey protein. Proximity of bait and prey proteins enables complementary DHFR fragments to assemble into functional DHFR, enabling growth on media containing methotrexate. (B) Proximity interactions are shown as ratio Z-scores calculated from the colony area of each GFP-prey-expressing strain with a full-length Vrl1 bait (Vrl1^FL^-DHFR^N^) divided by the colony area of an isogenic strain expressing the truncated Vrl1 control bait (Vrl1^ΔCt^-DHFR^N^) which cannot localize to endosomes. Significant proximity interactors (Z > 2) are shown in orange. The direct Vrl1 interactors Vin1 (Shortill et al., 2022) and Glc7 (Frier et al., 2026) are indicated. (C) The NAPA screen shares hits with the conventional DHFR screen (Shortill et al., 2022). (D) List of the top 12 hits from the nanobody-adapted DHFR screen. Coloured squares denote whether interactors are novel or were identified in the original screen (Shortill et al., 2022). (E) GO analysis of hits from the NAPA screen. The most specific term from each hierarchical subclass of enriched GO terms is shown as the log of the p value from a Bonferroni-corrected binomial test. Ontology terms also enriched in the original screen (Shortill et al., 2022) are indicated in dark green, and newly identified terms are shown in light green. (F) Vrl1 promotes endosomal localization of select Rab GTPases. Fluorescence micrographs of FM4-64-stained strains each expressing an endosomal Rab GTPase with an N-terminal GFP tag, and either *VRL1^FL^* or the truncated *VRL1^ΔCt^* allele whose product cannot localize to membranes. (G) Automated quantitation of GFP puncta per cell in *F*. Two-tailed unpaired t tests; n=3, cells/strain/replicate ≥ 1054; not significant, n.s.=p > 0.05, **=p < 0.01. Scale bars, 2 µm. Error bars report SEM. DHFR^N^, N-terminal fragment of dihydrofolate reductase. DHFR^C^, C-terminal fragment of dihydrofolate reductase. Nb, nanobody. FL, full-length. Ct, C-terminus. GO, gene ontology. Fold enrich., fold enrichment. OE, Overexpressed. NAPA, nanobody-adapted proximity assay. VINE, VPS9 GEF-interacting sorting nexin.

NAPA screening yielded 117 interactors with Z-scores of 2.0 or greater (Figure 1B and Supplementary Table 1) after normalizing to a truncated *VRL1^ΔCt^-DHFR^N^* control bait that lacks the phox homology-like (PX-like), BAR and ankyrin repeat-containing (AnkRD) domains required for VINE formation and association with endosomal membranes (Shortill et al., 2022). This list of putative Vrl1 interactors included 96 novel interactors and 21 of the 69 hits that were previously found in the DHFR^C^ screen (Figure 1C; Shortill et al., 2022). In addition to two previously identified partners of Vrl1, namely the obligate VINE subunit Vin1 (Z-score=32.6) and the Vrl1 AnkRD interactor Glc7 (Z-score=3.3; Frier et al., 2026), strong hits included multiple endosomal Rab GTPases and their effectors (Figure 1D). Gene ontology (GO) terms enriched among Vrl1 NAPA interactors included those found in the original interactome (Shortill et al. 2022) as well as novel terms such as vesicle tethering (Figure 1E; Ashburner et al., 2000; The Gene Ontology Consortium, 2026). The NAPA method therefore captures an extensive interactome that demonstrates VINE’s proximity to Rab GTPase signaling.

### VINE promotes Ypt10 localization to late endosomes

Among the top proximity interactors of Vrl1 were the Rab5-related GTPases Vps21, Ypt52, and Ypt10 (Figure 1D). Association of Rab5-family GTPases with endosomal membranes is stabilized by nucleotide exchange catalyzed by VPS9-family proteins (Cabrera and Ungermann, 2013). While extensive research has characterized Vps21 and Ypt52 function and addressed their regulation by VINE (Gerrard et al., 2000; Peplowska et al., 2007; Markgraf et al., 2009; Li et al., 2019; Frier et al., 2026), much less is known about Ypt10, which exhibits relatively weak localization to endosomes (Nesterova et al., 2026). To compare the effects of VINE on the localization of different endosomal Rab proteins, we observed each GFP-tagged Rab GTPase, as well as the Vrl1 binding partner Vin1, in strains expressing either full-length Vrl1 (Vrl1^FL^) or the non-functional Vrl1^ΔCt^ allele (Figure 1F, G and Figure S1A). Full-length Vrl1 decreased the number of bright GFP-Vps21 puncta per cell, whereas the number of bright GFP-Ypt53 and GFP-Ypt52 puncta was either unchanged or increased (Figure 1F, G), consistent with prior reports that VINE both promotes Rab5-family GTPase signaling and activates a Vps21-specific GAP (Bean et al., 2015; Frier et al., 2026).

Surprisingly, Vrl1^FL^ also caused GFP-Ypt10 to form bright puncta, suggesting VINE promotes Ypt10 localization to endosomes (Figure 1F, G and Figure S1B, C). Ypt10 has been previously reported to colocalize with endosomal markers in *VRL1*-deficient strains (Langemeyer et al., 2020; Nesterova et al., 2026; Louvet et al., 1999). Consistent with a previous report (Nesterova et al., 2026), GFP-tagged Ypt10 expressed from its native promoter in a *vrl1* mutant strain exhibited only dim GFP signal overlapping with endosomal mCherry-Vps21 puncta (Figure S1B).

The strikingly enhanced localization of Ypt10 upon *VRL1* expression suggests that Ypt10 signaling could be significantly impaired in the absence of VINE (Figure S1C). The increased endosomal targeting of Ypt10 is not simply due to increased VPS9-family GEF activity, because the overexpression of *VPS9*, which elevates the activity of other Rab5-family proteins (Frier et al., 2026), had no discernible effect on the localization of overexpressed, mCherry-tagged Ypt10 (Figure 2A, B). The prominent mCherry-Ypt10 puncta that appear upon *VRL1* expression (Figure 2A, B) therefore suggest that VINE plays a specialized role in Ypt10 activation. Indeed, *VRL1* expression caused an increase in GFP-Ypt10 puncta even in a strain lacking *MUK1* and *VPS9*, showing that VINE-dependent membrane targeting of Ypt10 does not require these other GEFs (Figure 2C, D). The strongly increased punctate localization of Ypt10 seen upon expression of *VRL1* suggested that VINE may function as a GEF for Ypt10. Indeed, the Vrl1^D373A^ allele, which has a mutation in the catalytic site of the VPS9 domain, was unable to target Ypt10 to puncta despite localizing to endosomes at comparable levels to wild-type Vrl1 (Figure 2E, F; Shortill et al., 2022). Together, these data support VINE as a Ypt10 GEF.

**Figure 2.**
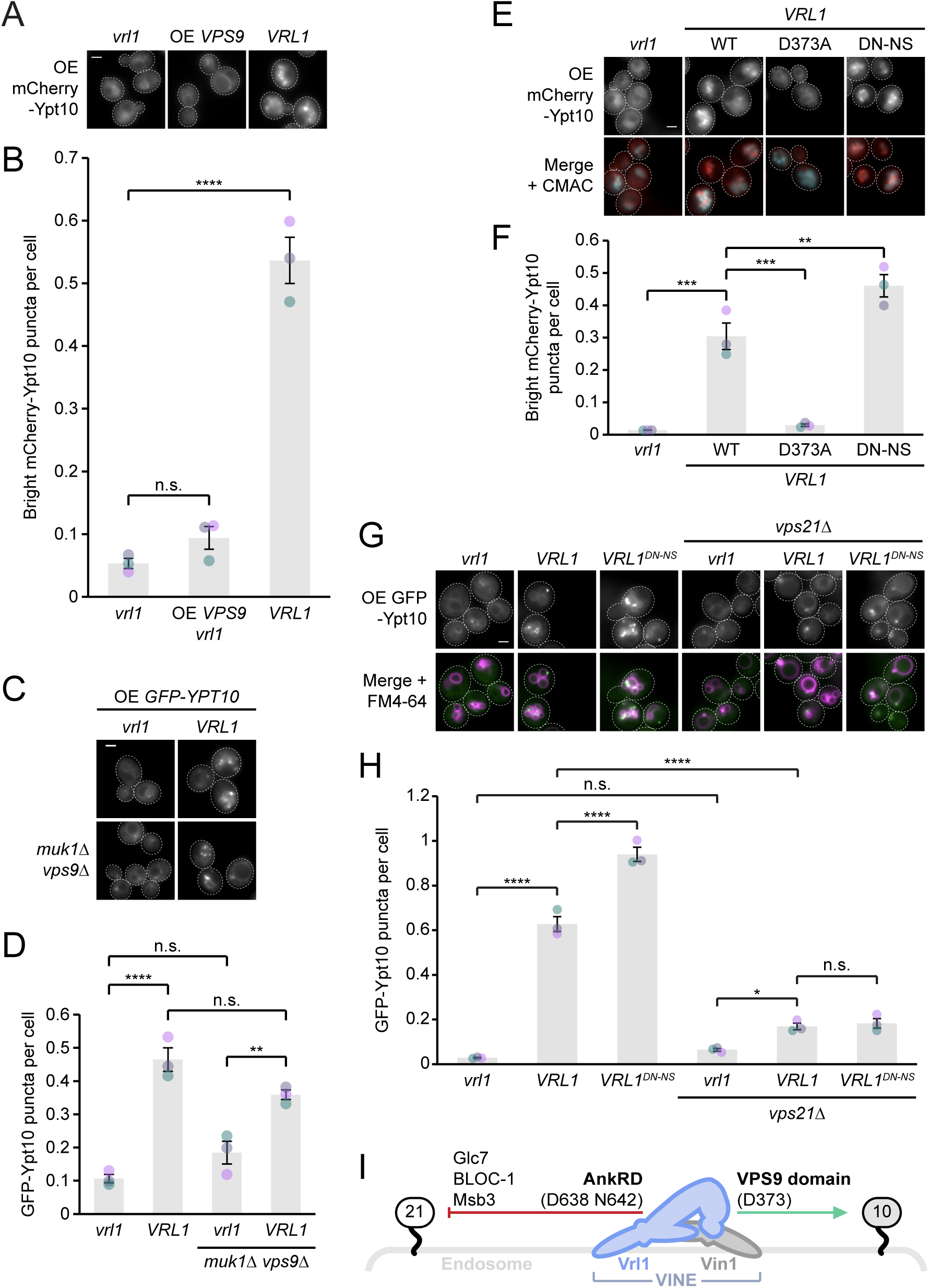
Ypt10 functions downstream of Vps21 and VINE. (A) Ypt10 is selectively targeted to endosomal puncta by Vrl1. Fluorescence micrographs of mCherry-Ypt10, showing effects of *VPS9* overexpression (OE) or *VRL1* expression. (B) Automated quantitation of mCherry-Ypt10 puncta per cell in *A*. Ordinary one-way ANOVA with Dunnett’s correction for multiple comparisons; n=3, cells/strain/replicate ≥ 1184; not significant, n.s.=p > 0.05, ****=p < 0.0001. (C) Other VPS9-family GEFs are not required for Ypt10 punctate localization. Fluorescence micrographs of GFP-Ypt10 with and without *VRL1* expression, showing effect of deleting other VPS9-family GEFs. (D) Automated quantitation of mCherry-Ypt10 puncta per cell in *C*. Ordinary one-way ANOVA with Tukey’s correction for multiple comparisons; n=3, cells/strain/replicate ≥ 663; not significant, n.s.=p > 0.05, **=p < 0.01, ****=p < 0.0001. (E) Vrl1 GEF activity promotes Ypt10 localization. Fluorescence micrographs of CMAC-stained cells expressing mCherry-Ypt10, showing effects of expression of *VRL1* separation-of-function alleles. (F) Automated quantitation of mCherry-Ypt10 puncta per cell in *E*. Ordinary one-way ANOVA with Dunnett’s correction for multiple comparisons; n=3, cells/strain/replicate ≥ 869; **=p < 0.01, ***=p < 0.001. (G) Ypt10 localizes downstream of Vps21. Fluorescence micrographs of FM4-64-stained cells expressing GFP-Ypt10, showing effects of Vrl1 expression and deletion of *VPS21*. (H) Automated quantitation of GFP-Ypt10 puncta per cell in *G*. Ordinary one-way ANOVA with Tukey’s correction for multiple comparisons; n=3, cells/strain/replicate ≥ 908; not significant, n.s.=p > 0.05, *=p < 0.05, ****=p < 0.0001. (I) Schematic of Rab regulation by VINE. Left: D638 and N642 residues in the Vrl1 AnkRD mediate Vps21 inactivation via Glc7, BLOC-1 and Msb3. Right: D373 in the VPS9 domain promotes Ypt10 signaling, likely through GEF activity. Scale bars, 2 µm. Error bars report SEM. OE, overexpressed. DN-NS, D638N N642S. VINE, VPS9 GEF-interacting sorting nexin.

We recently described a VINE-dependent pathway that culminates in GAP-dependent inactivation of the Rab5 homolog Vps21 (Frier et al., 2026). To determine whether Ypt10 signaling is influenced by this pathway, we used a separation-of-function mutation in the AnkRD of Vrl1 (*VRL1^D638N^ ^N642S^*, referred to as *VRL1^DN-NS^*) that blocks hyperactivation of the GAP Msb3 but leaves the GEF domain intact (Frier et al., 2026). Interestingly, the “GEF-only” Vrl1^DN-NS^ allele caused a slight increase in the number of bright mCherry-Ypt10 puncta per cell relative to wild type Vrl1 (Figure 2E, F). This suggests that the VINE-dependent increase in Msb3 GAP function not only inactivates Vps21 but also slightly dampens Ypt10 signaling.

One explanation for this is that VINE, while activating Ypt10 via its GEF domain, also promotes GAP activity on Ypt10 via its AnkRD. Alternatively, if Ypt10 functions downstream of Vps21, AnkRD-dependent regulation of Vps21 could indirectly affect Ypt10. We found that while deleting *YPT10* had no significant effect on GFP-Vps21 localization (Figure S2A, B), deleting *VPS21* drastically reduced the number of GFP-Ypt10 puncta per cell, and Ypt10 localization was severely impaired even with *VRL1* expression, suggesting Vps21 is required to establish a robust Ypt10 signaling domain at endosomes (Figure 2G, H). Importantly, Vps21 was not absolutely required for VINE to target Ypt10 to endosomes, as *VRL1* expression caused a slight but significant Vps21-independent increase in Ypt10 localization (Figure 2G, H). This suggests that VINE primarily acts downstream of Vps21 to promote Ypt10 signaling. Indeed, GFP-tagged Vrl1 also required *VPS21* for efficient endosomal localization (Figure S2C, D). This is expected, as a major determinant of VINE localization is the Vps21 effector PI 3-kinase (Shortill et al., 2022; Zhao et al., 2022; Špokaitė et al., 2026). Together these data suggest that Vps21 might promote endosomal targeting of Ypt10 through the PI3P-dependent recruitment of VINE.

Consistent with this model, mutations in the AnkRD that prevent VINE from promoting GAP function had no impact on Ypt10 in the absence of Vps21 (Figure 2G, H), indicating that the AnkRD-dependent GAP-activating pathway does not directly inactivate Ypt10. Together, these results suggest that while VINE acts through the Vrl1 AnkRD to promote Vps21 inactivation, VINE positively regulates Ypt10 through the Vrl1 VPS9 GEF domain (Figure 2I).

The finding that Vps21 signaling promotes the localization of VINE and thereby its substrate Ypt10 suggests a Rab cascade-like mechanism. We reasoned that if Vps21 acts upstream of Ypt10 in a cascade, the distribution of these two Rab proteins across endolysosomal membranes should be different. In agreement with a previous report that Ypt10 colocalizes with Vps21 (Nesterova et al., 2026), GFP-Ypt10 exhibited partial colocalization with mCherry-Vps21 when both Rabs were expressed at or near endogenous levels (Figure S1B). However, while Vps21 labeled both perivacuolar and peripheral populations of endosomes, Ypt10 puncta in Vrl1-expressing cells were almost exclusively perivacuolar, suggesting Ypt10 occupies mature endosomal membranes (Figure 3A, B; Hicke et al., 1997).

**Figure 3.**
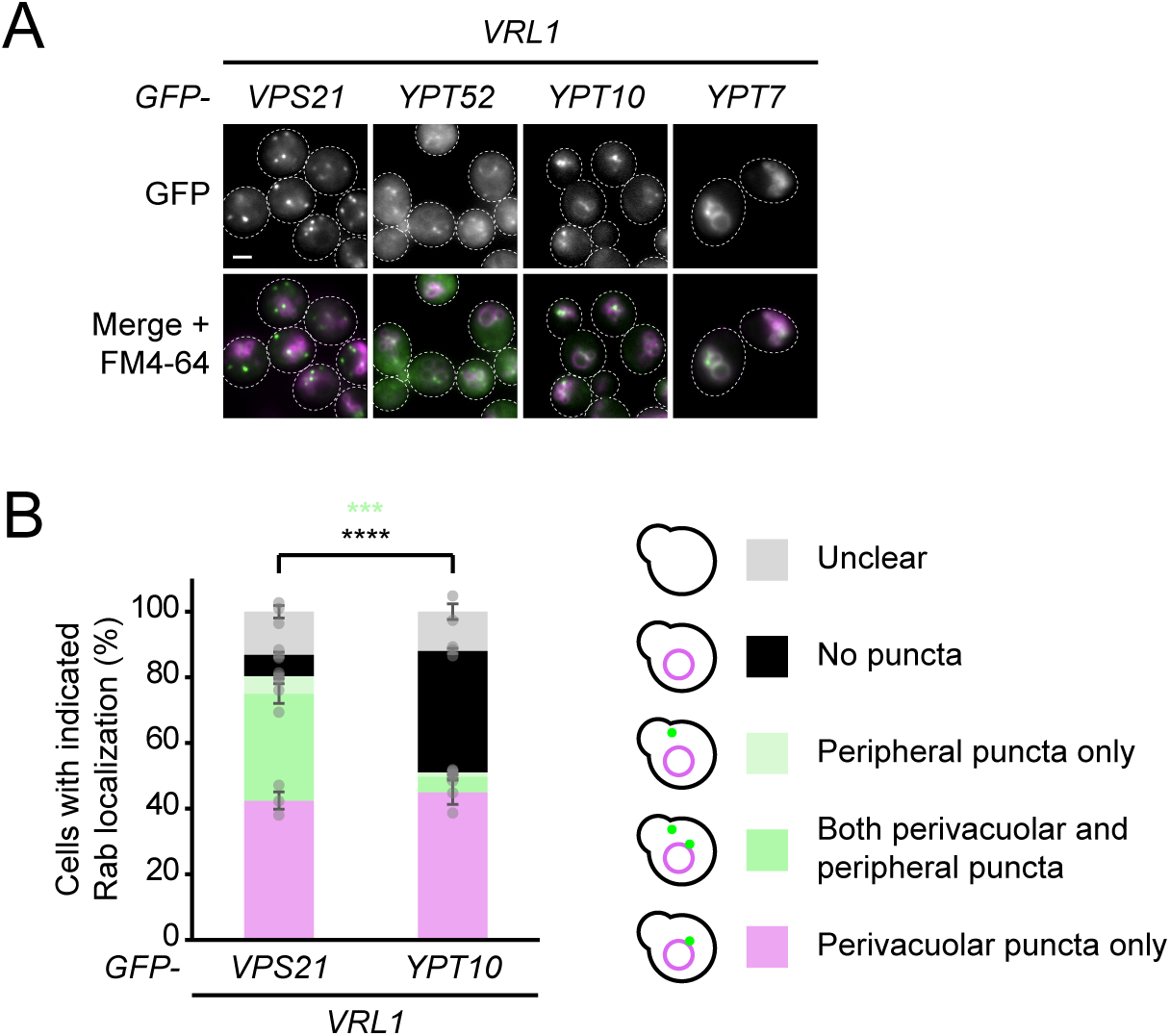
Ypt10 localizes to late endosomes. (A) Ypt10 localizes to perivacuolar endosomes. Fluorescence micrographs of GFP-tagged Rab GTPases in *VRL1*-expressing cells with FM4-64-stained vacuoles. (B) Position of GFP-Vps21 and GFP-Ypt10 puncta relative to vacuoles in *A*. Legend shows strategy for manual quantitation of blinded images. Two-tailed unpaired equal variance t test; n=3, cells/strain/replicate ≥ 324; ***=p < 0.001, ****=p < 0.0001. Result of each statistical test is shown in the colour corresponding to the localization pattern tested. Only significant differences are shown. Scale bars, 2 µm. Error bars report SEM.

Additionally, while Vps21 localized only to puncta, vacuolar membranes were populated not only by Ypt7 but also by Ypt52 and Ypt10 (Figure 3A). These results suggest that while Vps21 localizes to both peripheral early endosomes and perivacuolar late endosomes, Ypt10 signaling occurs specifically on late endosomes and can persist after endosome-to-vacuole fusion.

### Proximity interactomes of four endolysosomal Rab GTPases

Our data suggest that downstream of Vps21, VINE promotes Ypt10 signaling at late endosomes. This raised the possibility that a late endosomal pool of Ypt10 recruits effectors to promote a specific membrane trafficking event. We reasoned that a Ypt10 proximity interactome would include its functional partners and thus provide insight into Ypt10’s trafficking functions. Further, because our data indicate that VINE is critical for the robust localization of Ypt10 to late endosomes, these proximity interactions should be *VRL1*-dependent.

To find specific partners of Ypt10, we used the NAPA system to compare the proximity interactomes of Ypt10, Vps21, Ypt52 and Ypt7 (Figure 4A). We screened for interactors of each Rab GTPase bait in a strain lacking *VRL1* and in strains expressing *VRL1^WT^*, GEF-inactive *VRL1^D373A^*, or GEF-only *VRL^DN-NS^* which cannot induce Vps21 inactivation (Supplementary Tables 2-5). The overall interaction profile for Ypt10 was strikingly altered by the absence of functional *VRL1* or by a mutation in the GEF active site (Figure 4B, C). This is consistent with our microscopy data showing that membrane targeting of Ypt10 is strongly enhanced by Vrl1 GEF activity (Figure 2E, F). In contrast, the absence of functional *VRL1* or Vrl1 GEF activity had only a modest global effect on the Vps21 and Ypt52 interactomes, which is unsurprising given that these Rabs are also activated by Vps9 and Muk1 (Figure 4B). No *VRL1* allele substantially altered the interactome of Ypt7 (Figure 4B). Excluding the GDP dissociation inhibitor, Gdi1, which binds to Rab GTPases in their inactive GDP-bound form, all significant Ypt10 interactors were VINE-dependent. This further supports the conclusion that while other VPS9-family GEFs compensate for VINE loss to promote Vps21 and Ypt52 signaling, VINE is required for robust Ypt10 membrane targeting.

**Figure 4.**
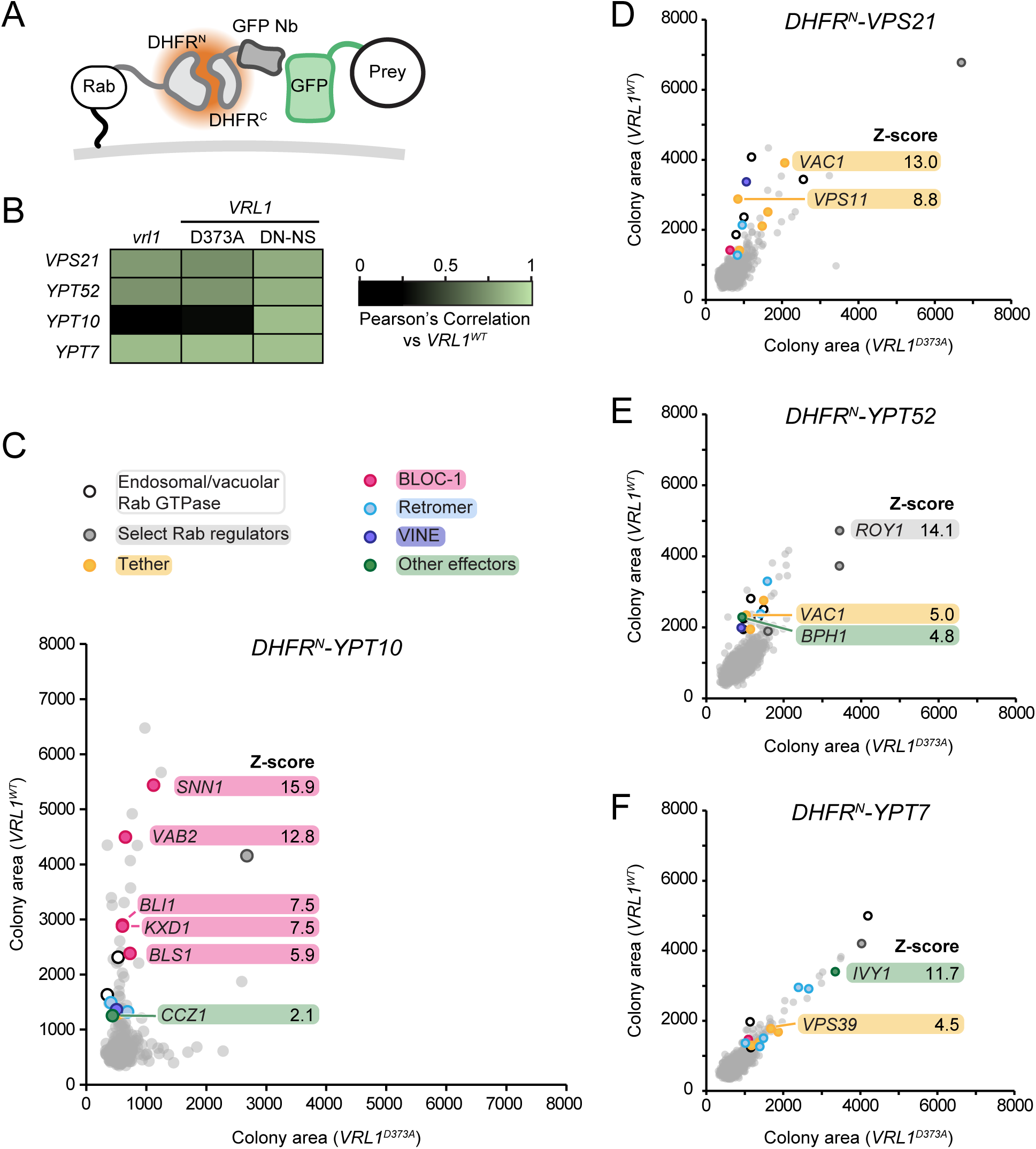
Ypt10 proximity interactors act in the endosomal Rab cascade. (A) Schematic of the nanobody-adapted proximity assay (NAPA). Each Rab GTPase bait is tagged at its N-terminus with the N-terminal fragment of DHFR (DHFR^N^). Prey proteins are GFP-tagged at their N-termini, and a GFP-binding nanobody (Nb) targets the C-terminal fragment of DHFR (DHFR^C^) to the prey protein. Proximity of bait and prey proteins enables complementary DHFR fragments to assemble into functional DHFR, enabling growth on media containing methotrexate. (B) Vrl1 GEF activity determines Ypt10 proximity interactions. Heatmap showing effects of *VRL1* mutations on genome-wide Z-score profiles for each Rab bait. For each genome-wide Z-score profile, the Pearson’s correlation to the Z-score profile for the same bait with *VRL1^WT^* is represented on a scale from black (no correlation) to green (perfect correlation). (C) Vrl1 GEF activity-dependent proximity interactors of DHFR^N^-Ypt10 include subunits of BLOC-1 and Mon1-Ccz1. Colony areas of strains expressing DHFR^N^-Ypt10, Nb-DHFR^C^ fusion and GFP-tagged preys with GEF-inactive *VRL1^D373A^*(x axis) are plotted against colony area of corresponding GFP prey strains expressing DHFR^N^-Ypt10, Nb-DHFR^C^ fusion and wild-type *VRL1* (y axis). Interactors of interest that were unique to Ypt10 are indicated. Legend shows categories of hits of interest and colour-coding scheme used in *C-F*. (D) Proximity interactors of DHFR^N^-Vps21 include multiple endosomal tethers. Data are presented as in *C*. (E) Proximity interactors of DHFR^N^-Ypt52 include the Ypt52-specific effector Bph1. Data are presented as in *C*. (F) Proximity interactors of DHFR^N^-Ypt7 include the Ypt7 effector Ivy1 and subunits of retromer and HOPS. Data are presented as in *C*. DHFR^N^, N-terminal fragment of dihydrofolate reductase. DHFR^C^, C-terminal fragment of dihydrofolate reductase. Nb, nanobody. WT, wild-type. DN-NS, D638N N642S.

Each Rab GTPase proximity interactome included hits consistent with previous reports of Rab-specific pathways (Figure S3A, Figure 4C-F). For example, a top Vps21 interactor was the tether Vac1, which specifically requires Vps21 for endosomal membrane targeting (Cabrera et al., 2013; Figure 4D), while the Ypt52 interactome included the Ypt52 regulator Roy1 (Liu et al., 2011) and Ypt52-specific effector Bph1 (Duarte et al., 2022; Figure 4E). Ypt7 effectors detected included all five subunits of retromer, the HOPS subunit Vps39, and Ivy1 (Seals et al., 2000; Balderhaar et al., 2010; Numrich et al., 2015; Figure 4F, Figure S3A,B). These results demonstrate that NAPA screening can capture Rab-specific interactions.

The Ypt10 interactome strikingly included five of six subunits of the endosomal biogenesis of lysosome-related organelles complex 1 (BLOC-1) as well as Ccz1, a subunit of the Ypt7 GEF Mon1-Ccz1 (Figure 4C, Figure S3A; Nordmann et al., 2010; John Peter et al., 2013). This is consistent with a previous report that Ypt10 and Mon1-Ccz1 exhibit strong binding *in vitro* and colocalize at endosomes (Langemeyer et al., 2020). BLOC-1 and Mon1-Ccz1 both function in the endosomal Rab cascade in which Vps21-positive membranes lose active Vps21 and acquire active Ypt7 (Rana et al., 2015; Borchers et al., 2021). These data further support the VINE-dependent late endosomal localization of Ypt10, and suggest that Ypt10 is present at sites of endosomal membrane maturation (Figure S3B).

### Ypt10 affects endosomal localization of potential effectors

This unique proximity of BLOC-1 and Ypt10 prompted us to ask whether Ypt10 can recruit BLOC-1 to membranes. BLOC-1 was previously reported to require Vps21 for its endosomal localization in *VRL1*-deficient laboratory yeast, indicating that in the absence of VINE, Ypt10 cannot support BLOC-1 localization (John Peter et al., 2013). We found that deleting *VPS21* severely reduced Bls1 localization even when *VRL1* was expressed to promote endosomal Ypt10 signaling (Figure 5A, B). However, because the membrane targeting of VINE and Ypt10 is severely impaired in a *vps21*Δ mutant (Figure 2G, H, Figure S2C, D), whether Ypt10 contributes to BLOC-1 recruitment downstream of Vps21 signaling remains to be determined.

**Figure 5.**
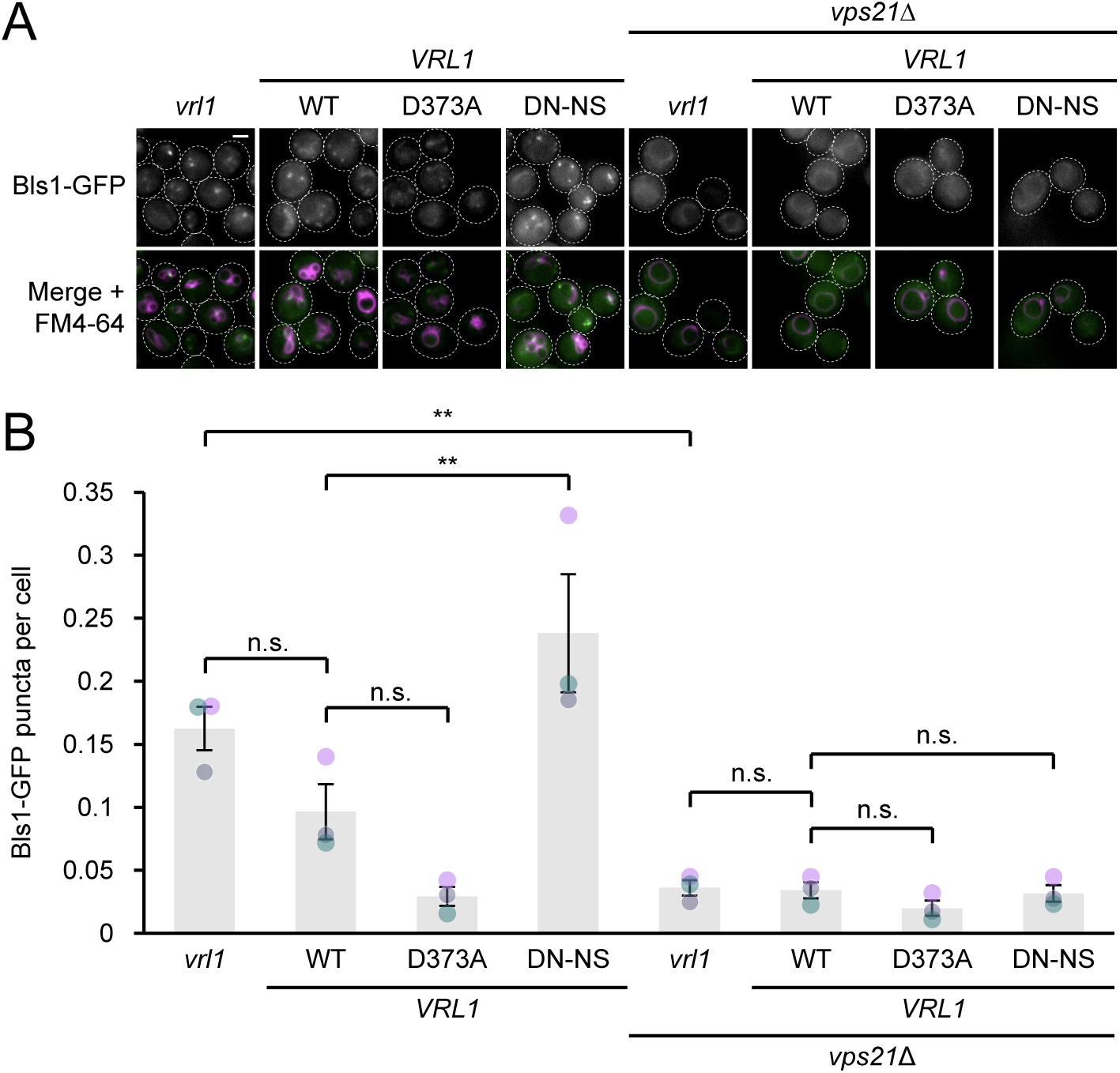
VINE-dependent Ypt10 signaling does not compensate for the loss of Vps21. (A) Vps21 is required for the endosomal recruitment of BLOC-1 and for effects of *VRL1*. Fluorescence micrographs of FM4-64-stained cells expressing the BLOC-1 subunit Bls1 as a fusion with GFP, showing effects of *VPS21* deletion and expression of *VRL1* separation-of-function alleles. (B) Automated quantitation of Bls1-GFP puncta per cell in *A*. Ordinary one-way ANOVA with Tukey’s multiple comparisons test; n=3, cells/strain/replicate ≥ 857; not significant, n.s.=p > 0.05, **=p < 0.01. Scale bars, 2 µm. Error bars report SEM. DN-NS, D638N N642S.

The Mon1-Ccz1 GEF complex represents a second candidate effector. Ypt10 has been reported to bind to Mon1-Ccz1 much more effectively than other endosomal Rab GTPases *in vitro* (Langemeyer et al., 2020) and our NAPA data suggest that VINE facilitates a specific proximity interaction between Ypt10 and Mon1-Ccz1 *in vivo* (Figure 4C). This suggests that upon its activation by VINE, Ypt10 may recruit Mon1-Ccz1 to late endosomes. While wild-type *VRL1* had no overall effect on the localization of Mon1 tagged with mScarletI (mScI), *VRL1* separation-of-function alleles revealed that Vrl1 GEF activity promoted endosomal localization of Mon1-mScI while AnkRD-dependent Vps21 inactivation opposed this (Figure 6A, B). Importantly, the “GEF only” allele *VRL1^DN-NS^* required *YPT10* to increase Mon1-mScI puncta, suggesting Ypt10 contributes to Mon1-Ccz1 recruitment in the presence of VINE (Figure 6C, D). These results support Mon1-Ccz1 as a potential effector of Ypt10.

**Figure 6.**
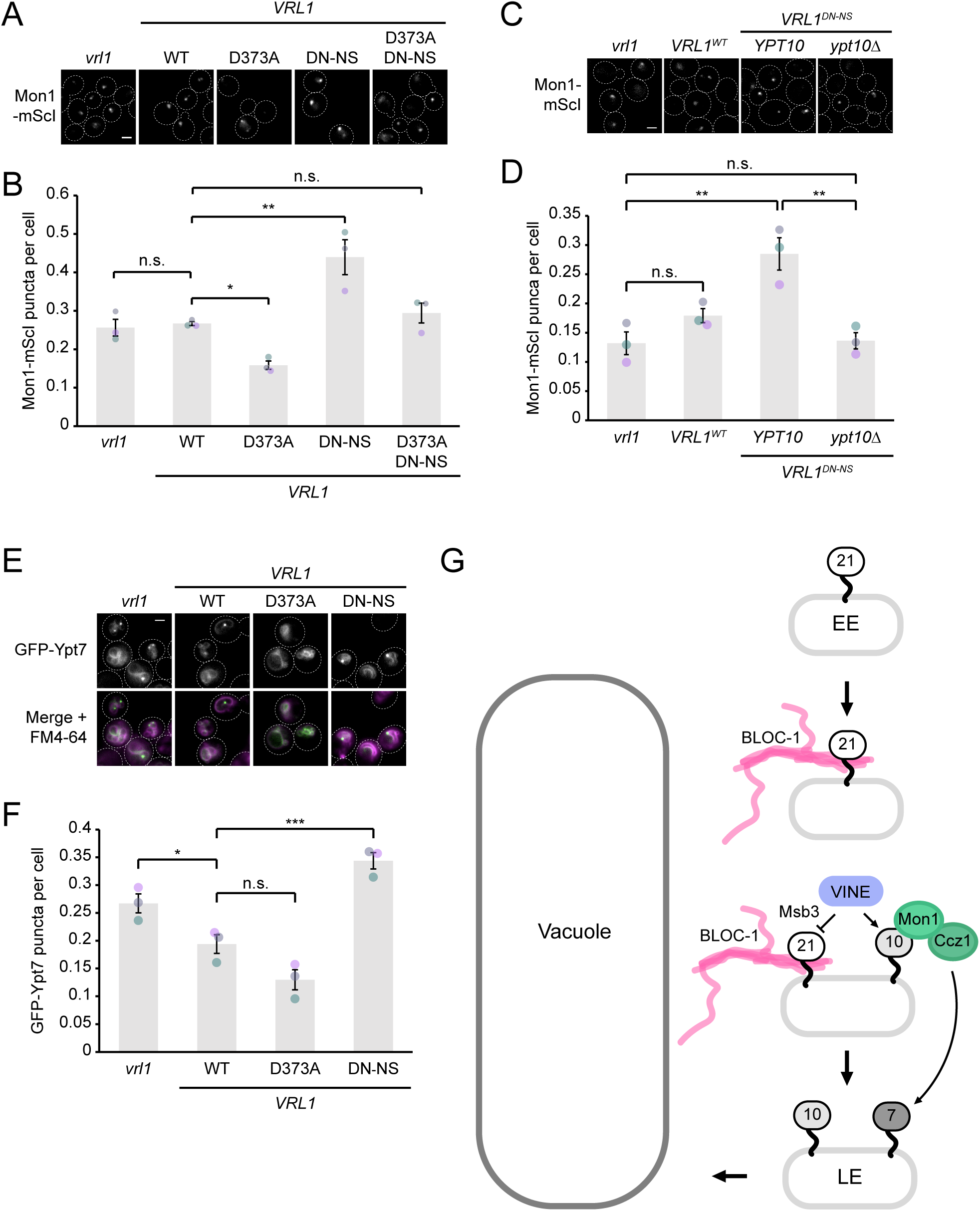
VINE-dependent Ypt10 signaling contributes to endosomal recruitment of the Ypt7 GEF Mon1-Ccz1. (A) Separate activities of VINE promote or suppress Mon1-Ccz1 endosomal localization. Fluorescence micrographs of Mon1-mScarletI (-mScI), showing the effects of expressing *VRL1* separation-of-function alleles. (B) Automated quantitation of Mon1-mScI puncta per cell in *A*. Ordinary one-way ANOVA with Dunnett’s multiple comparisons test; n=3, cells/strain/replicate ≥ 1147; not significant, n.s.=p > 0.05, *=p < 0.05, **=p < 0.01. (C) Increased endosomal Mon1-Ccz1 localization induced by the “GEF-only” allele of Vrl1 is *YPT10*-dependent. Fluorescence micrographs of Mon1-mScI, showing effects of wild-type *VRL1, VRL1^DN-NS^* which can activate Ypt10 without inactivating Vps21, and *YPT10* deletion. (D) Automated quantitation of Mon1-mScI puncta per cell in *C*. Ordinary one-way ANOVA with Tukey’s multiple comparisons test; n=3, cells/strain/replicate ≥ 2585; not significant, n.s.=p > 0.05, **=p < 0.01. (E) Rab regulation by VINE affects endosomal localization of Ypt7. Fluorescence micrographs of sfGFP-Ypt7 in FM4-64-stained cells expressing *VRL1* separation-of-function alleles. (F) Automated quantitation of sfGFP-Ypt7 puncta per cell in *E*. Ordinary one-way ANOVA with Dunnett’s multiple comparisons test; n=3, cells/strain/replicate ≥ 1530; not significant, n.s.=p > 0.05, *=p < 0.05, ***=p < 0.001. (G) Model for endosomal Rab regulation by VINE. Early endosomal membranes populated by Vps21 become proximal to vacuoles and acquire BLOC-1. VINE promotes both Ypt10 activation and Msb3-dependent Vps21 inactivation. The Ypt10 effector Mon1-Ccz1 activates Ypt7, leading to protein transport to the vacuole. Scale bars, 2 µm. Error bars report SEM. mScI, mScarletI. WT, wild-type. DN-NS, D638N N642S. EE, early endosome. LE, late endosome. BLOC-1, biogenesis of lysosome-related organelles complex 1. VINE, VPS9 GEF-interacting sorting nexin.

### Rab regulation by VINE fine-tunes endosomal Ypt7 signaling

Mon1-Ccz1 activates Ypt7 at late endosomes to facilitate endosome-to-vacuole transport (Cabrera et al., 2009; Nordmann et al., 2010). The contributions of VINE and Ypt10 to Mon1-Ccz1 recruitment raised the possibility that Ypt10 could promote Ypt7 activation at endosomal membranes. In this model, VINE both accelerates Vps21 inactivation and establishes a late endosomal pool of Ypt10 to promote Mon1-Ccz1 activity, leading to Ypt7 activation. If VINE thus aids a transition from Vps21 to Ypt7 signaling, it could prime the endosomal membrane for anterograde trafficking to the vacuole.

Based on this model, VINE should enhance the endosomal pool of Ypt7. Indeed, when comparing *VRL1* separation-of-function alleles, the “GEF-only” *VRL^DN-NS^* allele increased Ypt7 localization at endosomes relative to WT *VRL1*(Figure 6E, F). However, VINE-mediated inactivation of Vps21 appears to counter this GEF-dependent effect and limit Ypt7 recruitment, likely by reducing the endosomal localization of Ypt10 and Mon1-Ccz1 (Figure 2E, F and Figure 6A, B). Under steady-state, non-stress conditions, these opposing activities of VINE are balanced, resulting in a small overall reduction in endosomal Ypt7. We hypothesize that regulatory events could influence the equilibrium between these VINE activities to increase the endosomal pool of Ypt7 and drive the Rab cascade.

These results contribute new details to a model of Rab GTPase regulation during endosomal maturation (Figure 6G). In this model, early endosomes populated by Vps21 mature, acquiring BLOC-1 and becoming more proximal to vacuoles. At late endosomes, endosomal Rabs recruit Mon1-Ccz1 to activate Ypt7 and facilitate protein transport to the vacuole. We propose that by inactivating Vps21 and activating Ypt10, VINE could exclude active Vps21 from Ypt7 signaling domains.

## Discussion

Here we identify VINE as a positive regulator of the small GTPase Ypt10. Our results suggest VINE acts as a GEF for Ypt10 to promote Ypt10 localization at late endosomal membranes. VINE-dependent Ypt10 activation increased the proximity of Ypt10 to potential effectors including Mon1-Ccz1, which is a known Ypt10 binding partner and the GEF for the downstream GTPase Ypt7 (Nordmann et al., 2010; Langemeyer et al., 2020). Because VINE also suppresses the related GTPase Vps21 (Frier et al., 2026), VINE could replace an endosomal pool of Vps21 with Ypt10, leading to increased Ypt7 activity and priming the late endosomal membrane for Ypt7-dependent fusion with the vacuole (Wang et al., 2002; Cabrera et al., 2009; Balderhaar et al., 2010).

### An auxiliary module in the late endosomal Rab cascade

Detecting Ypt10 among the strongest proximity interactors of VINE was unexpected, as endosomal targeting of Ypt10 is relatively weak (Langemeyer et al., 2020). Indeed, we observed poor endosomal localization of Ypt10 in strains lacking VINE, suggesting Ypt10 may have little or no activity at endosomes in the absence of VINE. This could explain why identifying clear *in vivo* functions for Ypt10 has been challenging in studies of VINE-deficient yeast.

Ypt10 activation is not the only known role of VINE in Rab GTPase regulation, as we recently found that VINE promotes Vps21 inactivation via the GAP Msb3 (Frier et al., 2026). We propose that these regulatory events are coordinated to support protein trafficking as part of a broader Rab regulatory program. VINE’s GEF activity toward Ypt10, coupled with its GAP-mediated inactivation of Vps21, suggests that VINE may coordinate a regulated transition in Rab identity known as Rab switching.

The late endosomal Rab cascade, which replaces active Rab5 with a Rab7 signaling domain (Rink et al., 2005), is a well-characterized example of Rab switching. The recruitment of the Rab7 GEF complex Mon1-Ccz1 by a Rab5-family GTPase (Nordmann et al., 2010; Langemeyer et al., 2020; Borchers et al., 2023), followed by Rab7-coupled Rab5 inactivation, is essential for endosomal maturation (Chotard et al., 2010; Poteryaev et al., 2010; Rana et al., 2015; Tu et al., 2022; Ott et al., 2025). Our findings suggests that VINE adds an intermediate step to the analogous cascade in yeast: by coupling Vps21 inactivation to Ypt10 activation, it replaces active Vps21 with Ypt10 during membrane maturation. The subsequent interaction between Ypt10 and the Ypt7 GEF Mon1-Ccz1 (Langemeyer et al., 2020) would then mediate the transition to Ypt7 signaling.

Because the late endosomal Rab cascade has been observed in *S. cerevisiae* strains that lack VINE, the VINE-Ypt10 module is not essential for recruiting Mon1-Ccz1 and promoting Ypt7 activity (Langemeyer et al., 2020). Similarly, the subtle trafficking phenotypes upon *YPT10* deletion suggest that without active Ypt10, other Rabs such as Vps21 are sufficient to recruit Mon1-Ccz1 (Louvet et al., 1999; Nesterova et al., 2026). If VINE-dependent Ypt10 is not essential for endosomal maturation, what is its role? Vps21 additionally recruits endosomal tethers to promote fusion of endosomal membranes and vesicular carriers, thus a close involvement of Vps21 in Ypt7 activation might inappropriately introduce unsorted proteins during the transfer of sorted anterograde cargo to the vacuole (Russell et al., 2012; Cabrera et al., 2013). Replacing Vps21 with Ypt10 could allow Mon1-Ccz1 to be recruited while preventing such interference. VINE and Ypt10 may therefore represent an intermediate step in the Vps21-to-Ypt7 Rab cascade that insulates Vps21 and Ypt7 signaling domains and supports high-fidelity cargo trafficking to the vacuole.

Consistent with the model that VINE activates Ypt10 for a specialized role in Mon1-Ccz1 recruitment, we found that VINE GEF activity and Ypt10 promote the localization of Mon1 at endosomes. However, this effect was opposed by the VINE-dependent inactivation of Vps21, resulting in a balanced effect on endosomal levels of Mon1 and Ypt7. While such zero-sum regulation appears counterintuitive, it may provide an opportunity to regulate Ypt7 activation at endosomes: signaling events could tip this balance, allowing VINE to accelerate or delay Ypt7 signaling. In fact, Rab7 activation and endosomal traffic are influenced by multiple inputs (Cabrera et al., 2009; Lawrence et al., 2014; Yasuda et al., 2016; Langemeyer et al., 2020), and both the membrane recruitment of Mon1-Ccz1 and its activation are independently regulated (Borchers et al., 2023). These findings point to strict control of Ypt7 function at endosomes. Identifying regulators of VINE’s GEF or GAP-promoting activities could therefore reveal additional inputs controlling the rate of Ypt7 activation and endosome-to-vacuole transport.

### Alternative models of Ypt10 function

Although our data support a role for Ypt10 in the late endosomal Rab cascade, Ypt10 may have additional functions in membrane trafficking. Rab switching is increasingly recognized as a widespread feature of vesicular trafficking, and transport vesicles must inactivate the Rab from the source membrane and activate another Rab to facilitate fusion at the target membrane. Indeed, membrane coats formed by retromer can recruit both GAPs and GEFs including the Vrl1 homolog VARP, enabling Rab activation and inactivation to be coupled to the dynamic process of retrograde trafficking (Hesketh et al., 2014; Jia et al., 2016; Antón-Plágaro et al., 2025). Thus, VINE could couple Ypt10 activation to Vps21 inactivation during a process such as the formation of tubular transport carriers. The homology of the VINE subunit Vin1 to a retromer component suggests VINE could itself be a trafficking coat as well as a Rab regulator (Shortill et al., 2024). VINE contains a BAR dimer and is therefore predicted to activate Ypt10 at sites of membrane curvature, such as retrograde sorting domains (Kovtun et al., 2018; Shortill et al., 2022). If VINE is found to oligomerize and form membrane tubules analogous to retromer, Vps21 inactivation could prevent fusion with the source membrane while Ypt10 activation could promote interactions with unknown effectors to mediate cargo delivery to a target membrane.

Identifying additional Ypt10 effectors will clarify its functions throughout the endolysosomal system. Further investigation of the striking proximity between Ypt10 and BLOC-1 may reveal shared functions of these proteins. Although VINE promotes Ypt10 targeting to late endosomes, other GEFs can direct Ypt10 to the cell periphery to support endocytic trafficking independently of VINE (Nesterova et al., 2026). These findings suggest that different GEFs may activate Ypt10 at distinct membranes in the endolysosomal pathway. Variation in membrane lipid composition likely contributes to compartment-specific GEF activity by influencing both GEF function and the membrane association of Rab proteins (Kulakowski et al., 2018; Cezanne et al., 2020; Klink et al., 2022). Future work may uncover whether these endocytic and late endosomal pools of Ypt10 share effectors and functions.

The physiological importance of VINE and Ypt10 remains unclear, as the loss of VINE and late endosomal Ypt10 in laboratory settings produces only subtle phenotypes in the endolysosomal system. A possible explanation is that this GEF-Rab module provides specialized or inducible regulation of endosomal protein trafficking. The membrane trafficking system plays an essential role in enabling cells to respond rapidly to external stressors, and most environmental conditions encountered by wild yeast are not easily reproduced in the laboratory. VINE-dependent Ypt10 activity may improve fitness in natural environments while being dispensable under culture conditions that are optimal for yeast survival. Further study of Ypt10 and VINE, especially under conditions that stress the endolysosomal system, promises a more complete understanding of endosomal maturation and protein trafficking.

## Materials and Methods

### Yeast strains and plasmids

Information regarding yeast strains and plasmids used in this study can be found in Tables S6 and S7 respectively. Sequences of oligonucleotides used in this study can be found in Table S8. Yeast strains were constructed using homologous recombination, with alterations to genomic DNA confirmed using colony PCR and microscopy as appropriate. Plasmids were constructed using Golden Gate cloning or NEBuilder (New England BioLabs, Ipswich, Massachusetts) in *Escherichia coli* or by homologous recombination in yeast and confirmed by DNA sequencing. As all yeast strains used contain an inactive *vrl1* allele, expression of functional VINE was induced by introducing plasmids bearing functional *VRL1* alleles.

### Split-DHFR protein fragment complementation assay

A MATα strain expressing both a DHFR^N^-tagged bait and a fusion of DHFR^C^ with the GFP-binding nanobody LaG16 (Fridy et al., 2014) was mated to the MATa SWAT collection of N-terminally GFP-tagged preys (Weill et al., 2018). After selection of diploid yeast, a BM3-BC robot (S&P Robotics Inc., Toronto, Canada) was used to replicate array to media containing methotrexate. CellProfiler (Lamprecht et al., 2007) was used to measure colony area after four days at 30°C. For the Vrl1-DHFR^N^ screen, the area measurement of each Vrl1^FL^-DHFR^N^-expressing colony was divided by the area of the corresponding Vrl1^ΔCt^-DHFR^N^-expressing colony, and ratios used to calculate Z-scores in Microsoft Excel 2025 (Microsoft, Redmond, Washington). For Rab-DHFR^N^ screens, the nonspecific hit *NPT1* was removed before colony area values were used to calculate Z-scores. Each bait was screened asynchronously. The Gene Ontology (Ashburner et al., 2000; Gene Ontology Consortium, 2021) GO enrichment analysis tool (Thomas et al., 2022) was used to identify significantly enriched terms in each DHFR proximity interaction dataset. To generate heatmaps summarizing DHFR interaction data, Microsoft Excel 2025 (Microsoft, Redmond, Washington) was used to calculate the Pearson correlation coefficient (*r*) between colony area values for two baits within an experiment and to place *r* values on a colour scale ranging from black (0 ≤ r ≤ 0.25) to green (*r* = 1.0).

### Fluorescence microscopy and automated image analysis

Yeast strains were cultured in minimal selective media to log phase and added to glass-bottom 96-well plates (Cellvis, Mountain View, California) treated with concanavalin A. When appropriate, vacuoles were visualized by staining with 100 μM 7-amino-4-chloromethylcoumarin (CMAC) (Setareh Biotech, San Jose, California) or 4 μM FM4-64 (Invitrogen, Waltham, Massachusetts), which were incubated with yeast for 30 minutes prior to washing and a 60-minute incubation at 30°C prior to imaging. Widefield images were taken using a Leica DMi8 microscope (Leica Microsystems, Wetzlar, Germany) with ORCA-flash 4.0 digital camera (Hamamatsu Photonics, Shizuoka, Japan) and Plan-Apochromat 63x/1.30 glycerol immersion lens (Leica Microsystems, Wetzlar, Germany). MetaMorph 7.8.13.0 (MDS Analytical Technologies, Sunnyvale, California) was used for image acquisition. All intensity scaling adjustments were linear and consistent across all images of a given fluorophore within an experiment. Figures were constructed in Illustrator CC 2021 (Adobe, San Jose, California) using images imported via Photoshop CC 2025, with the application of bicubic expansion where appropriate (Adobe, San Jose, California). MetaMorph 7.8.13.0 journals (MDS Analytical Technologies, Sunnyvale, California) were used for automated quantitation of images. Count Nuclei was used to mask dead cells and identify live cells, and Granularity was used to identify punctate fluorescence signal based on feature size and intensity above local background in unmodified images. ImageJ (Schneider et al., 2012) Multi-Point tool was used for manual quantitation of uniformly scaled blinded images.

### Statistical analysis and graphical representation of quantitative data

Microsoft Excel 2025 (Microsoft, Redmond, Washington) was used to generate graphs of quantitative data. Error bars show standard error of the mean. GraphPad Prism 10.2.2 (GraphPad Software, San Diego, California) was used to conduct statistical tests. Normality of data was assumed but not formally tested. Hypotheses were tested against a 95% confidence threshold (P < 0.05).

## Supplemental Material

Figures S1-S2 contain the results of fluorescence microscopy experiments. Tables S1-S5 contain proximity interaction data for Vrl1 and Rab GTPase baits. Figure S3 contains additional comparisons and analyses of results from four split-DHFR screens measuring Rab proximity interactomes. Tables S6-S8 respectively detail the yeast strains, plasmids and oligonucleotides used in this study.

## Data Availability Statement

All data generated in this study are included in the manuscript and supplemental files.

## Acknowledgements

We thank Dr. Maya Schuldiner (Weizmann Institute of Science, Rehovot, Israel) for sharing the SWAT yeast library. This work was supported by the Natural Sciences and Engineering Research Council of Canada (grants RGPIN-2022-04573 and RGPIN-2016-04290 to EC); Canada Foundation for Innovation (Leading Edge Fund 30636); Canadian Institutes for Health Research (grant PJT-180544 to EC, CGS-M and CGS-D Frederick Banting and Charles Best Canada Graduate Scholarships to MSF); and the University of British Columbia (Affiliated Fellowships and 4-Year Doctoral Fellowship to MSF).

## Abbreviations

AnkRD: ankyrin repeat-containing domain
BAR: Bin-amphiphysin-Rvs
BLOC-1: biogenesis of lysosome-related organelles complex 1
Ct: C-terminus
DHFR: dihydrofolate reductase
DN-NS: D638N N642S
FL: full-length
GAP: GTPase activating protein
GEF: guanine nucleotide exchange factor
mScI: mScarletI. NAPA: Nanobody-adapted proximity assay
VINE: VPS9 GEF-interacting sorting nexin.

## Supplementary Figure and File Legends

**Supplementary Figure 1.**
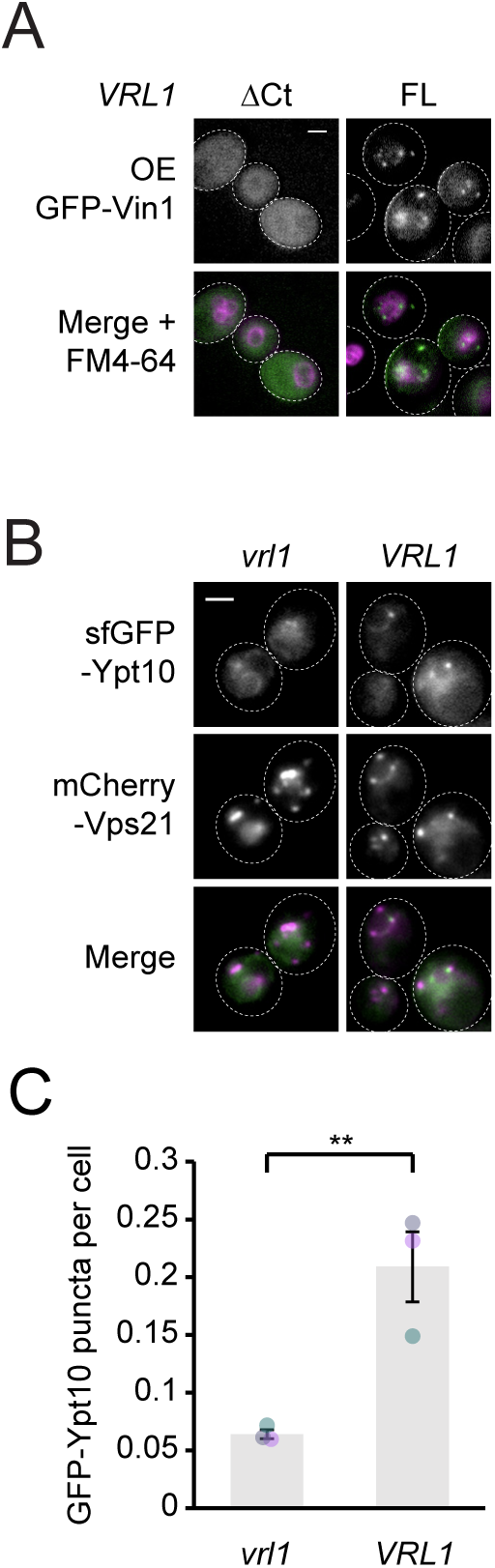
(A) Full-length Vrl1, but not C-terminally truncated Vrl1, enables Vin1 localization to endosomes. Fluorescence micrographs of FM4-64-stained cells expressing GFP-Vin1 from the *NOP1* promoter and expressing either full-length Vrl1 or Vrl1^ΔCt^. (B) VINE promotes endosomal localization of Ypt10 expressed at endogenous levels. Fluorescence micrographs of cells expressing sfGFP-Ypt10 from its native promoter and expressing the endosomal marker mCherry-Vps21 from the moderate *RNR2* promoter. (C) Automated quantitation of sfGFP-Ypt10 puncta in *B*. Two-tailed unpaired equal variance t test; n=3, cells/strain/replicate ≥ 858; **=p < 0.01. Scale bars, 2 µm. Error bars report SEM. sfGFP, superfolder GFP. FL, full-length. OE, overexpressed. VINE, VPS9 GEF-interacting sorting nexin.

**Supplementary Figure 2.**
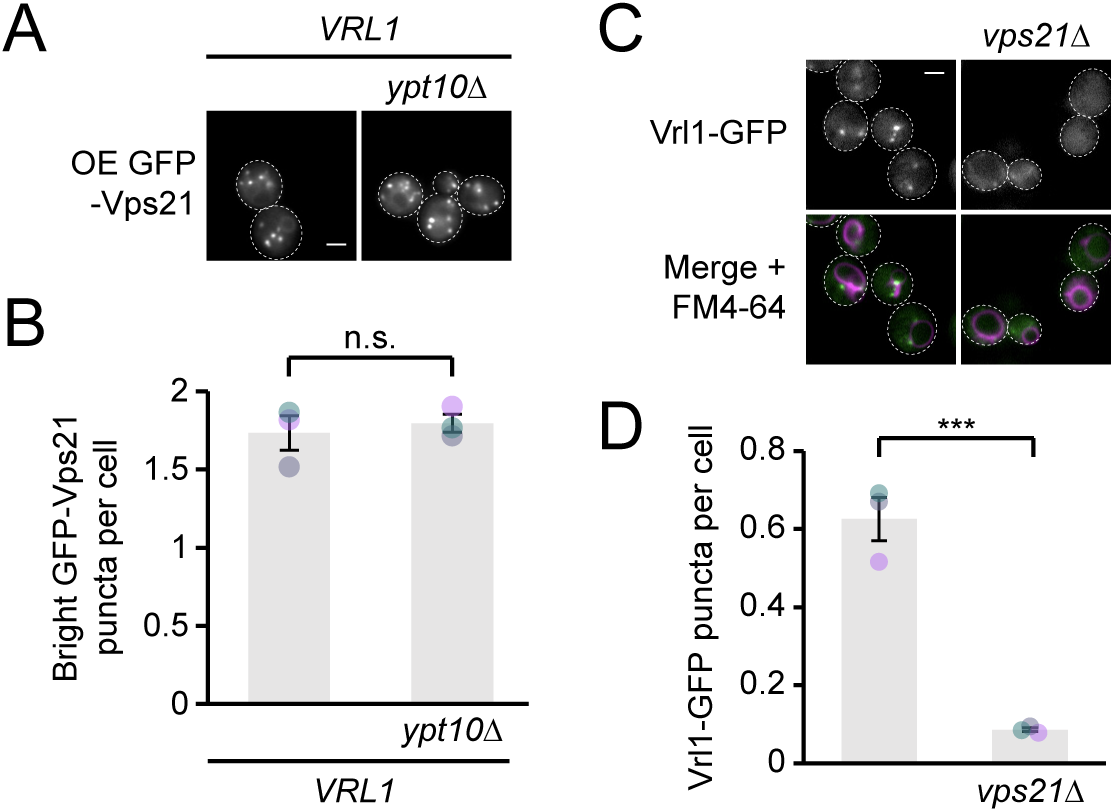
(A) Ypt10 does not influence Vps21 localization. Fluorescence micrographs of GFP-Vps21 in *VRL1*-expressing cells with or without a deletion of *YPT10*. (B) Automated quantitation of GFP-Vps21 puncta per cell in *A*. Two-tailed unpaired equal variance t test; n=3, cells/strain/replicate ≥ 542; not significant, n.s. = p > 0.05. (C) Endosomal localization of VINE requires Vps21. Fluorescence micrographs of FM4-64-stained cells expressing GFP-tagged Vrl1, showing effect of deleting *VPS21*. (D) Automated quantitation of Vrl1-GFP puncta in *C*. Two-tailed unpaired equal variance t test; n=3, cells/strain/replicate ≥ 1293; *** = p < 0.001. Scale bars, 2 µm. Error bars report SEM. VINE, VPS9 GEF-interacting sorting nexin.

**Supplementary Figure 3.**
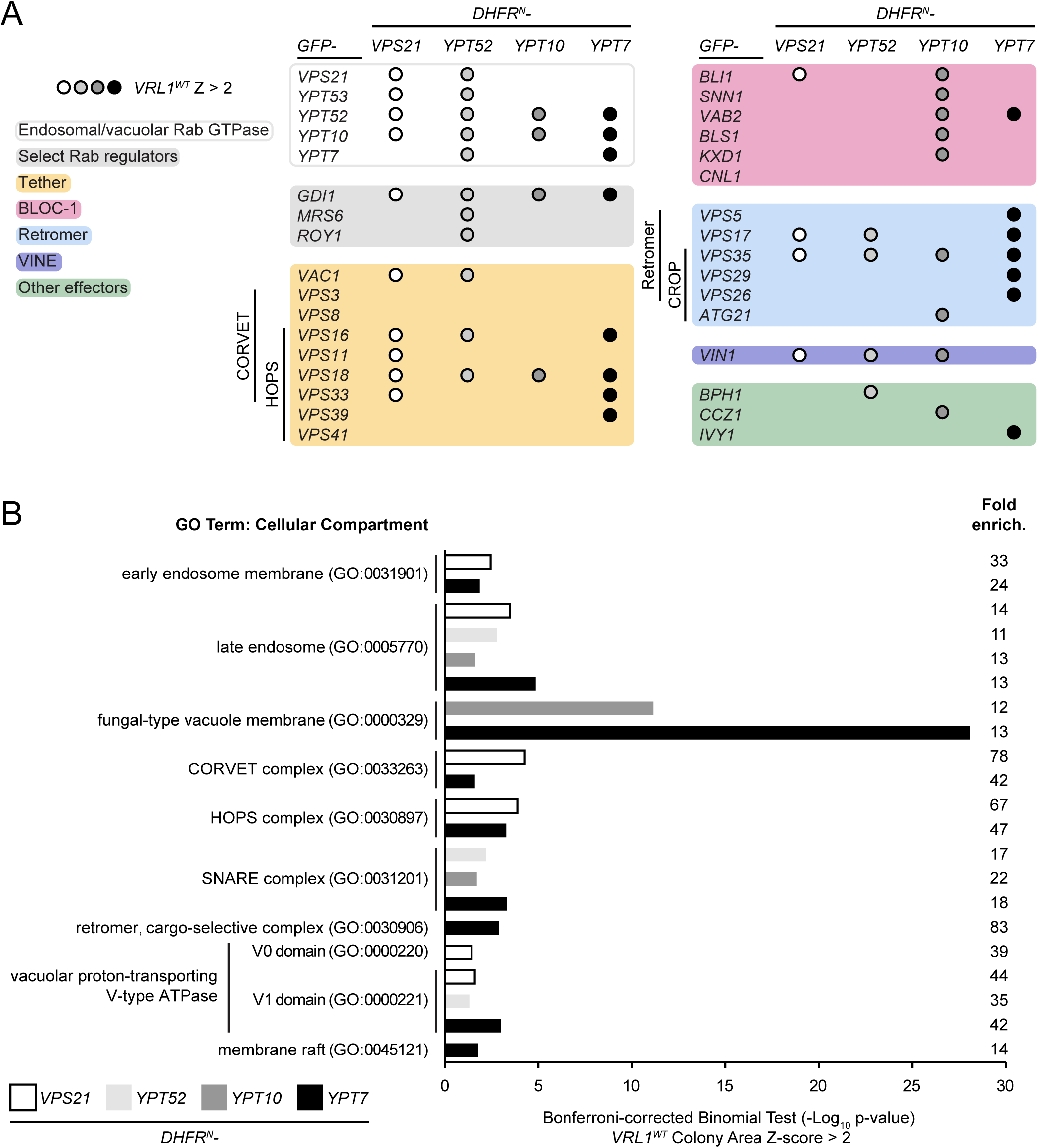
(A) Proximity interactions between Rab baits and all hits of interest in genome-wide DHFR screens. The presence of a circle indicates a significant interaction (Z>2) between the Rab GTPase bait and indicated prey. Each shade corresponds to a different Rab protein. (B) Gene ontology (GO) enrichment analysis of significant interactors from each split-DHFR screen. GO terms of the most specific hierarchical subclass with a fold enrichment value >10 are presented as the negative base 10 log of the associated p-value from a Bonferroni-corrected binomial test of significance. WT, wild-type. DHFR^N^, N-terminal fragment of dihydrofolate reductase. Fold enrich., fold enrichment.

**Table S1. Vrl1 proximity interactors.** Vrl1 FL/ΔCt ratio Z scores for all significant DHFR interactors (Z ≥ 2).

**Table S2. Ypt10 proximity interactors.** Colony area and Z-score values for all significant DHFR interactors (Z ≥ 2).

**Table S3. Vps21 proximity interactors.** Colony area and Z-score values for all significant DHFR interactors (Z ≥ 2).

**Table S4. Ypt52 proximity interactors.** Colony area and Z-score values for all significant DHFR interactors (Z ≥ 2).

**Table S5. Ypt7 proximity interactors.** Colony area and Z-score values for all significant DHFR interactors (Z ≥ 2).

**Table S6. List of *Saccharomyces cerevisiae* strains used in this study**

**Table S7. List of plasmids used in this study.**

**Table S8. List of oligonucleotides used in this study.**

